# A novel anti-influenza combined therapy assessed by single cell RNA-sequencing

**DOI:** 10.1101/2021.07.27.453967

**Authors:** Chiara Medaglia, Ilya Kolpakov, Yong Zhu, Samuel Constant, Song Huang, Arnaud Charles-Antoine Zwygart, Valeria Cagno, Emmanouil T. Dermitzakis, Francesco Stellacci, Ioannis Xenarios, Caroline Tapparel

## Abstract

Influenza makes millions of people ill every year, placing a large burden on the healthcare system and the economy. To develop a novel treatment against influenza, we combined virucidal sialylated cyclodextrins with interferon lambda and demonstrated, in human airway epithelia, that the two compounds inhibit the replication of a clinical H1N1 strain more efficiently when administered together rather than alone. We investigated the mechanism of action of the combined treatment by single cell RNA sequencing analysis and found that both the single and combined treatments impair viral replication to different extents across distinct epithelial cell types. We also showed that each cell type comprises multiple sub-types, whose proportions are altered by H1N1 infection, and assess the ability of the treatments to restore them. To the best of our knowledge this is the first study investigating the effectiveness of an antiviral therapy by transcriptomic studies at the single cell level.

## 1. Introduction

Influenza is a highly contagious respiratory infection that accounts every year for about ∼ 3 to 5 million cases of severe illness and up to 650,000 deaths (*1*). More than a century after the “Spanish” pandemic, the health systems are still struggling to cope with seasonal influenza, something that bodes poorly in the event of a novel pandemic. Influenza is caused, in humans, by influenza A (IAV) and influenza B (IBV) viruses. Although the latter are almost exclusively found in the human population, IAVs emerge from a huge zoonotic reservoir (*2*). In a process called antigenic drift IAVs rapidly acquire adaptive mutations allowing them not only to evade the host immune response but also to neutralize annual attempts to generate effective vaccines (*3*). As a consequence, seasonal epidemics endanger every year children, elderly people, pregnant women and people of any age with comorbid illnesses (*4*). In addition, due to their ability to cross the species barrier, IAVs pose a high pandemic risk. The arrangement of the viral genome on multiple RNA segments allows for exchange of genetic material between different viral strains which co-infect the same host, giving rise to novel gene-reassorted variants. This process, when accompanied by the expression of new surface glycoproteins, is named antigenic shift, as it results in the emergence of strains which infect immunologically naive humans and cause potentially pandemic outbreaks (*5*). Lastly, when the viral reassortants possess new virulence factors, they can be associated with increased pathogenicity.

Influenza virus (IV) is enveloped, with a negative single strand RNA genome. The viral protein hemagglutinin (HA) of human IV binds preferentially α2,6- linked sialic acid (Sia) moieties located on the surface of the host cell, thus triggering viral entry through clathrin-mediated endocytosis (*6*). Upon entering a new host, IV establishes an infection in the epithelial cells lining the upper airways (*7*). When the infection stays restricted to this region of the respiratory tract it causes a rather mild disease. But if it spreads to the lungs it can determine viral pneumonia, with progression to acute respiratory distress syndrome (ARDS) and death from respiratory failure (*8*). IAV disrupts the functions of the respiratory barrier by inducing epithelial cell death via intrinsic viral pathogenicity, or through a robust immune response (*9*). This alteration leads to exposure of new attachment sites for bacteria (*10*), thus making the host more vulnerable to secondary infections by other pathogens, which significantly contribute to the morbidity of influenza (*11*).

Annual vaccination is the cornerstone of prevention against IVs. However, the vaccine has to be adapted yearly and does not always match with circulating strains. This is further complicated by the co-circulation of different IV types and different IAV subtypes (*12*). In the 2017 to 2018 United States season, vaccine effectiveness was estimated to be only ∼25% against influenza A subtype H3N2 viruses, which however comprised ∼69% of infections (*13*). Antivirals represent an important second line of defense against IV, but all the currently available drugs are only efficient if taken at the early stages of the disease. Moreover, they inevitably exert selective pressure on the virus, which causes the appearance of drug-resistant variants (*14–16*). It results that there is an unmet need to develop novel therapies against IV. Several studies indicate IFN λ as a promising therapeutic candidate to control influenza and other viral respiratory diseases (*17, 18*). The family of IFN λ (alias IFN type III) comprises IFN λ1, IFN λ2 and IFN λ3 (also known as IL-29, IL-28A and IL-28B, respectively) and the recently identified IFN λ4 (*19*). Like IFN type I, IFN λ acts in an autocrine and in paracrine fashion, inducing an antiviral state through the expression of interferon-stimulated genes (ISGs), that inhibit viral replication at multiple steps (*20*). The distinct tract of IFN λ is a circumscribed range of action, as the expression of its receptor is mostly restricted to the epithelial cell surfaces (*21*). Indeed, immune cells are largely unresponsive to IFN λ (*21, 22*). Thus, while IFN type I targets nearly all immune cells, creating massive inflammation that may further weaken the host (*23*), IFN λ only acts at the epithelial barriers and on few innate immune cells, without causing immunopathology (*18, 24*). These properties suggest IFN λ as a treatment of choice against acute viral infections, such as influenza, with a higher tolerability than IFN type I. IFN λ plays a critical early role, not shared by IFN type I, in protection of the lung following IV infection (*25–28*) and several *in vivo* studies show that it also exerts variable degrees of antiviral activity against both IAV and IBV strains (*29*). It has been reported that, in B6.A2G-MX1 mice infected with H1N1 IAV, IFN λ intranasal administration prevents viral spread from the upper to the lower airway, without noxious inflammatory side effects (*26, 30*). Importantly human pegylated IFN λ1 passed both phase I and II clinical trials for hepatitis C treatment, displaying an attractive pharmacological profile (*31, 32*).

Combination therapy is considered a valuable approach to provide greater clinical benefit, especially to those at risk of severe disease. Combining drugs targeting different mechanisms of viral replication may increase the success rate of the treatment (*33, 34*), as also demonstrated in our previous work, showing that IFN λ1 co-administration delays the emergence of H1N1 IAV resistance to oseltamivir (*35*). We recently developed 6’SLN-CD [heptakis-(6-deoxy-6-thioundec)-beta-cyclodextrin grafted with 6’SLN(Neu5Ac-a-(2-6)-Gal-b-(1-4)-GlcNAc;6’-N-Acetylneuraminyl-N-acetyllactosamine](*36*) a non-toxic anti-influenza antiviral designed to target and irreversibly inactivate extracellular IV particles, preventing their entrance into the host cell. 6’SLN-CD significantly decreases IAV replication in both *ex vivo* and *in vivo* models of infection (*36*). However, 6’SLN-CD targets the globular head of IV HA, which undergoes constant antigenic drift, thus posing a concrete problem of resistance emergence [14a]. In this work we chose to combine human IFN λ1, the host frontline defense against IAV, with 6’SLN-CD, in order to increase its effect and lower the chances of antiviral resistance. The two compounds hinder viral replication on different fronts: IFN λ1 boosts the host innate response while 6’SLN-CD traps and inactivates newly formed virions. To mimic the *in vivo* environment, we assessed the combinatorial effect of the compounds in 3D human airway epithelia (HAE) reconstituted at the air-liquid interface (*11, 37*) and showed that IFN λ1 enhances 6’SLN-CD antiviral activity. HAE perfectly mimic both the pseudostratified architecture of the human respiratory epithelium, composed of basal, ciliated, and secretory cells, and its defense mechanisms. In addition, they allow the use of clinical viral specimens thus preserving their original pathogenicity and biological characteristics, which are inevitably lost upon repeated passages in cell lines (*37–40*).

As host cellular heterogeneity strongly impacts virus-host interplay and is mirrored in the response to antiviral treatments (*41, 42*), we investigated the mechanism of action of IFN λ1 plus 6’SLN-CD by single cell RNA sequencing (scRNA-seq). This approach allowed us to trace the landscape of the modifications through which individual cells respond to IAV infection and to the treatments. We found that in different epithelial cell types, both the individual and the combined antivirals hinder viral replication to different extents, depending on the permissiveness of the cells to H1N1. We also showed that each basal, secretory and ciliated cells comprise multiples subclusters, whose proportions are altered by the infection. Surprisingly even though in each cell type the antivirals reduced viral replication synergistically, they were not able to restore the changes in cell subcluster composition in a similar manner. Lastly, in absence of infection, IFN λ1 + 6’SLN-CD did not alter the proportions of the main epithelial cell types, further supporting the therapeutic potential of the formulation. The findings presented in this work pave the way to future *in vivo* experiments, to better assess the efficacy of IFN λ1 + 6’SLN-CD treatment against influenza. To the best of our knowledge this is the first study investigating the effectiveness of an antiviral treatment by scRNA-seq.

## 2. Results

### 2.1 IFN λ1 and 6’SLN-CD display synergistic activity against H1N1 IAV *ex vivo*

We optimized an IFN λ1/6’SLN-CD formulation to inhibit IAV in *ex vivo* 3D HAE. The tissues were infected with a clinical A/Switzerland/3076/2016 H1N1strain that has not been passaged in cell lines, to exclude any *in vitro* adaptation bias. First, we determined the best administration mode of the individual treatments. While 6’SLN-CD successfully inhibited viral replication when administered at 8 hours post infection (hpi) on the apical surface of the HAE, IFN λ1 reduced viral spread only when administered at 24h before infection (hbi), and on the basal side of the tissue (**Figure S1**). Even though IFN λ pre-treatment is not an ideal clinical option, our data are in line with already published *in vitro* and *ex vivo* studies, confirming the effectiveness of IFN λ only in pre-treatment and on the basal side of polarized epithelial tissues (*43, 44*). The underlying reason for that is the mechanism of action of IFN λ and its kinetic. Unlike 6’SLN-CD, which directly targets the virus and inactivate it within minutes (*36*), the antiviral state induced by IFN λ relies on the activation of a gene expression program which takes several hours to be effective. Of note, in mouse models of infection, IFN λ prevents IV spread when administered via the intranasal route in therapeutic use, i.e. once the clinical symptoms of the disease are manifested, which would correspond to an administration at the apical side in our settings (*26, 30*). The discrepancy of IFN λ antiviral effects between *in vivo* and *ex vivo* systems is due to the lack of immune cells in the latter. *In vivo*, IFN λ is sensed by the transepithelial dendritic cells of the respiratory mucosa, which strongly amplify its signal (*45*).

Based on these observations and to achieve the maximum combinatorial antiviral effect, we administered 6’SLN-CD and IFN λ1 according to the following protocol: HAE were first treated on their basal side with IFN λ1, starting at 24 hbi, while 6’SLN-CD was administered at 8 hpi on the apical side of the tissue. Both IFN λ1 and 6’SLN-CD were then co-administered daily up to 72 hpi (**Figure 1A**). The quantification of viral replication at both 48 and 72 hpi, by measuring the viral particles released from the apical surface of the tissues, revealed that when administered in combination IFN λ1 and 6’SLN-CD were more effective than when administered individually. The synergistic effect was evident at 48 hpi (≥ 1 log reduction for both individual treatments vs > 2 log reduction for the combined one) and it persisted at 72 hpi, when the antiviral effect of the individual treatments was lost (**Figure 1B**). Of note, IFN λ1 and 6’SLN-CD are non-toxic nor as individual (*32, 36*), nor as combined treatments (**Figure S2**). These data indicate that IFN λ1 treatment potentiates the antiviral action of 6’SLN-CD.

**Figure 1.**
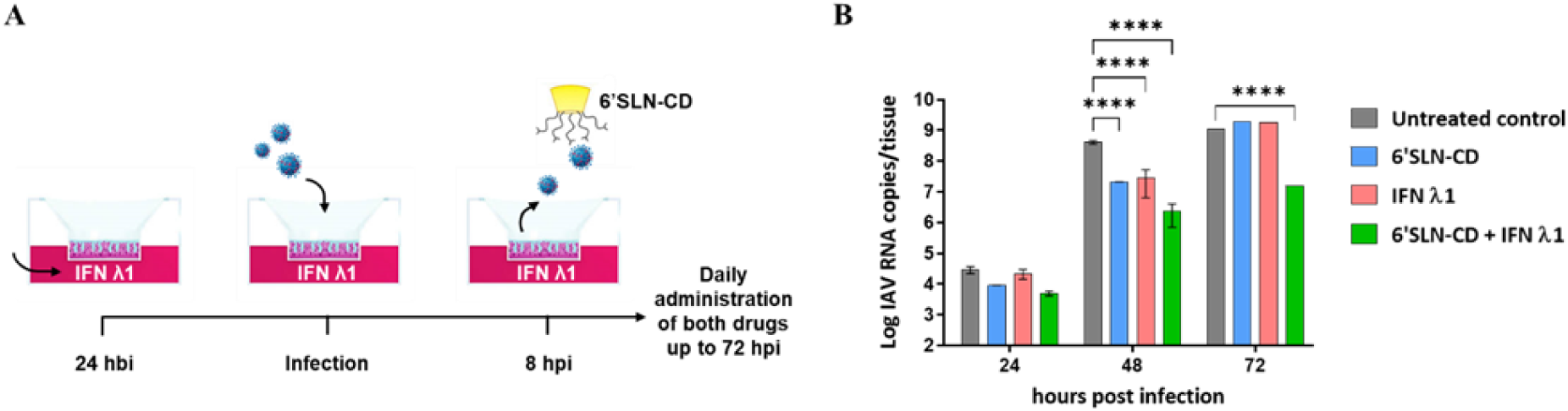
Combinatorial effect of 6’SLN-CD + IFN λ1 in HAE. **A)** Schematics of 6’SLN-CD and IFN λ1 (60 µg and 5.5 ng per tissue, respectively) combined administration. **B)** Bar plot showing the kinetic of IAV replication in HAE treated with 6’SLN-CD only, or with IFN λ1 only, or with both compounds according to A). The results represent three independent experiments conducted in duplicate in HAE developed from a pool of donors and infected with 10^3^ RNA copies of clinical A/Switzerland/3076/2016 H1N1 (0 h corresponds to the time of viral inoculation). Viral replication was assessed measuring the apical release of IAV by RT-qPCR. ***, *p* ≤ 0.001; ****, *p* ≤ 0.0001.

### 2.2 scRNA-seq analysis reveals that the proportions of HAE basal, secretory and ciliated cells are not affected by IAV infection, nor by the antiviral treatments

In order to investigate at the molecular level the mechanism of action of IFN λ1 + 6’SLN-CD and its effects on HAE, we performed scRNA-seq analysis on both non-infected and infected tissues, administered or not with the individual or combined treatments. When conducting transcriptomic studies, it is essential to reach a fair compromise between viral and host gene expression. Viral replication occurs at the expenses of the host transcription machinery, resulting eventually in a complete host shutoff (*46*). Preventing the expression of cellular proteins at multiple steps is also a strategy adopted by the virus to counteract the antiviral response (*47*). We selected the time of 48 hpi as the most suitable to perform scRNA-seq in our acute infection model, as it provides a wide window of analysis of both viral and host genes. At 48 hpi viral replication is in the exponential phase (**Figure 1B** and (*36*)) resulting, however, in a still low cytopathic effect (*11*) and in ∼ 10% infected cells, measured based on the expression of IAV nucleoprotein (NP) (**Figure S3**). Moreover, at this time point the advantage of the combined treatment is evident while individual treatments are still efficient, allowing to compare the therapeutic approaches with each other (**Figure 1B**). When correlating within the same HAE model, the number of IAV RNA copies measured from the apically released virus with the number of infected cells measured by FACS, we observed that the majority of the virus was produced by a small percentage of infected cells (**Figure S3 and 1B**). This finding is in line with previously published reports showing that between cells, there is a high level of variability in the outcome of IAV infection, which results from multiple sources of heterogeneity, such as the number of viral transcripts per cell, the antiviral response and the timing of the infection (*48, 49*).

sc-RNAseq relies on tissue dissociation, which can dramatically impact cell viability in epithelial tissues, as their survival is highly dependent on physical connections and communication between cells (*50, 51*). We established a dissociation protocol that allows to retrieve every cell type of the HAE (secretory, basal and ciliated cells, **Figure S4**) without compromising cell viability, thus preserving the quality of the mRNA within individual cells. To perform scRNA-seq analysis, HAE were infected with IAV and treated or not with 6’SLN-CD, IFN λ1, or with IFN λ1 + 6’SLN-CD. To assess the perturbations induced by the formulation in absence of virus, an uninfected control (mock) untreated and one treated with IFN λ1 + 6’SLN-CD were included. Cells were partitioned for cDNA synthesis and barcoded using the Chromium controller system (10x Genomics), followed by library preparation and sequencing (Illumina). Sample demultiplexing, barcode processing and gene counting was performed using the Cell Ranger analysis software (*52*). Following the inspection of standard quality-control metrics, we selected barcodes having > 10.000 reads and < 15% of mitochondrial reads. The selection procedure resulted in 12.778 captured cells (∼ 2.129 cells per condition). Of note, since our partitioning input was 4.000 cells per condition, the recovery rate was about 50%, which is in line with previously reported works (*52*).

The upper respiratory epithelium comprises several specialized cell types that likely respond to IAV infection in distinct ways (*53*). Using Seurat analytical pipeline, we performed an unsupervised graph-based clustering (*54*) on the Cell Ranger integrated dataset, comprising all the tested experimental conditions (**Figure 2A-C**). To match the identified clusters with the cell types found in the respiratory epithelium we used both cluster-specific and canonical marker genes (*55*) (**Figure 2B, D and S5A, B**).

**Figure 2.**
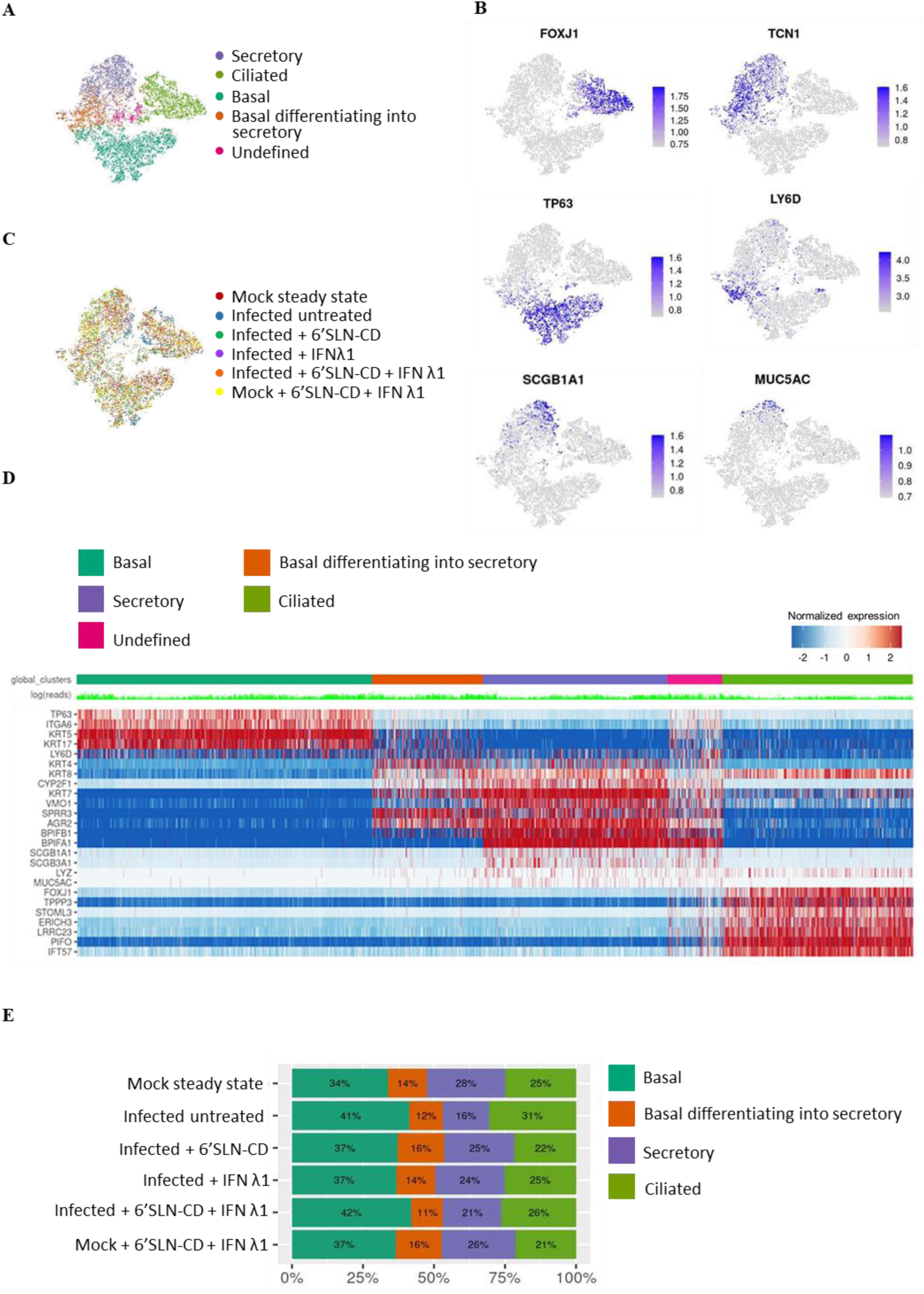
Analysis of HAE cell diversity. **A)** t-distributed stochastic neighbor embedding (t-SNE) visualization of the major cell types composing human HAE. Individual cell types were annotated using a combination of graph-based clustering results from Seurat and expression analysis of several canonical cell type-specific markers. The t-SNE plots shown in panels A-C are presented in the same spatial orientation (i.e. the location of cells expressing the canonical markers in figure B corresponds to the location of the specific cell types in panel A). **B)** t-SNE plots illustrating in blue the expression patterns of some of the canonical markers used to annotate the three main airway epithelial cell types: *FOXJ1* for ciliated cells, *TCN1* for all secretory cells, *TP63* for basal cells, *LY6D* for differentiating cells, *SCGB1A1* for club cells and MUC5AC for goblet cells; scale bars are Log2. **C)** t-SNE visualization of the scRNA-seq data for all single cells in the following conditions: mock steady sate, infected untreated (infected with A/Switzerland/3076/2016 H1N1strain), infected + 6’SLN-CD, infected + IFN λ1, infected + IFN λ1 + 6’SLN-CD, and mock + IFN λ1 + 6’SLN-CD. **D)** Heatmap representing the gene expression profiles of 12.778 single cells from human HAE grouped into five clusters. Expression values are Pearson residuals from SCTransform binomial regression model [70] fitted to UMI counts (see Methods). The cells were clustered solely on the expression of the shown hallmark genes. **E)** bar graph showing the relative percentage of each main epithelial cell type described above in each experimental condition described in C).

In all analyzed HAE we identified five distinct clusters. Three of them corresponded to mature basal (*TP63+/ITGA6+/KRT5* ^high^*/KRT17* ^high^), ciliated (*FOXJ1+/PIFO+/TPPP3+*) and to a mixed population of secretory cells, including both goblet-mucous (*MUC5AC+*) and club cells (*SCGB1A1+/SCGB3A1+*) **(Figure 2A-D)**. One cluster was made of a population of basal cells uniquely defined by high levels of *LY6D*, marker of cellular plasticity and differentiation (*56*) **(Figure 2A-D)**. Like *in vivo*, also *ex vivo* HAE basal cells have both self-renewing and multipotent properties (*57, 58*). The current consensus is that in steady state conditions, basal cells differentiate first into secretory cells that in turn give rise to ciliated cells (*59*). However, after injury, ciliated cells can be directly generated by basal cells (*59, 60*). *LY6D* ^high^ basal cells were characterized by the co-expression of both basal and secretory hallmark genes, such as *KRT5*, *KRT17*, and *BPIFB1*, *SPRR3*, *AGR2* respectively (**Figure 2D**). This cluster was hence identified as constituted by basal cells differentiating into secretory cells. The last cluster, consisting of 843 cells (6.6 % of the total selected cells), did not display a unique gene signature compared to the others, but co-expressed basal, secretory and ciliated hallmark genes (**Figure 2D and S5B**). It was also marked by an increased number of gene counts in comparison to the other clusters (**Figure S5C**). We therefore concluded that this cluster likely resulted from doublets and excluded it from further analysis. These observed cell types and their proportions (**Figure 2E**) are consistent with previous scRNA-seq studies and indicate that our *ex vivo* model recapitulates the respiratory epithelium *in vivo* (*61*).

We next determined the relative abundances of ciliated, secretory, basal and differentiating basal cells across different conditions (**Figure 2E**). We found that nor the infection alone, nor the treatments in the presence or absence of the infection, induced substantial changes in the relative proportions of these main epithelial cell types (**Figure 2E**). IAV causes a strong cytopathic effect which results in a significant loss of ciliated cells and important alterations of the tissue structure. However, in our HAE infection model this phenomenon occurs only at 120 hpi and it is therefore not evident at 48 hpi (*11*), which explains our results.

### 2.3 The antiviral treatments affect IAV replication to different extents across different HAE cell types

We next asked how viral transcripts would distribute across cell type clusters, in each experimental condition. Global analysis of both host and viral transcriptomes in all 11.935 cells revealed that at 48 hpi and in absence of treatments, IAV transcripts were detected in all cell types and were more abundant in ciliated cells, followed by secretory cells, basal differentiating into secretory and lastly, by basal cells (**Figure 3A** and **S6**). Basal cells are located in the lower part of the epithelium, do not reach the apical side and are therefore physically protected from the virus in the first stages of the infection, when the ciliated cell layer is preserved (*37*). Secretory cells have been shown to be the immediate target of IAV (*11, 62*), while ciliated cells become preferentially infected at later stages of infection (*63, 64*). Nonetheless, we asked whether the different numbers of viral transcripts between secretory and ciliated cells relied also on the expression levels of host factors involved in IV infection. We measured in steady state conditions the average mean expression of twelve cellular genes promoting multiple steps of IV replication (*65*), in secretory (including basal differentiating into secretory) vs ciliated cells (**Figure S7**). We found that secretory cells express higher levels of genes involved in IV RNA replication, such as CD151 (*66*) and HMGB1 (*67*), or in viral maturation and release like TMPRSS4 (*67*) and Rack1 (*68*). While ciliated cells express higher levels of CLTA(*69*) and EPS8 (*70*), necessary for viral endocytosis and uncoating (**Figure S7**). These data better explain the higher susceptibility to IAV infection of ciliated over secretory cells.

**Figure 3.**
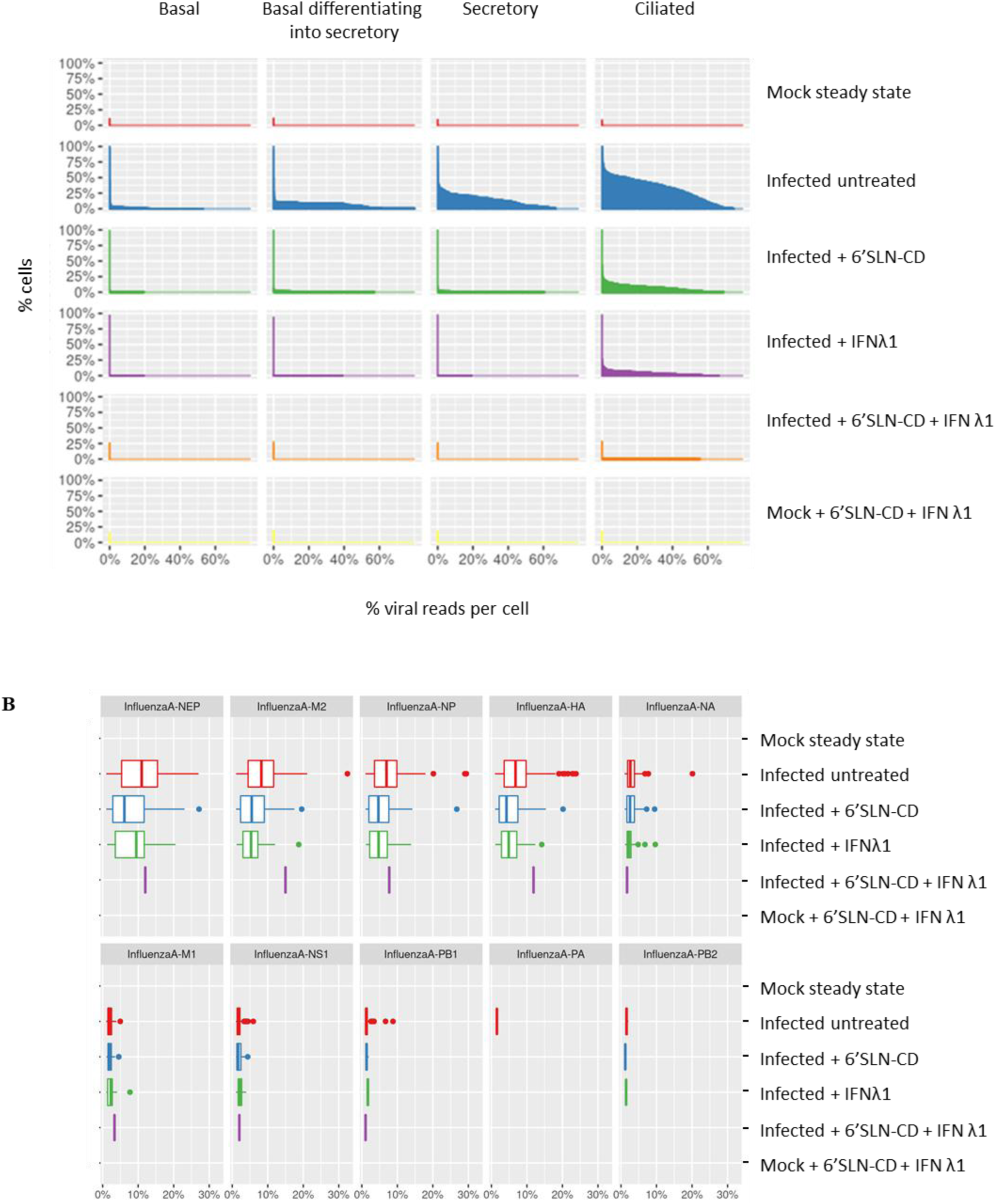
**A)** Distribution of IAV transcripts across cells having more than 1% of viral reads in different HAE cells clusters and across different experimental conditions. **B)** Box plot summarizing the relative fraction of viral mRNA for each IAV gene segment across all experimental conditions. Conditions as specified in figure 2C.

Compared to the mock steady state, in the 6’SLN-CD alone condition all main epithelial cell types displayed a decreased number of viral transcripts. Even so, this reduction was more pronounced in secretory (∼ 11-fold reduction) and in basal cells differentiating into secretory cells (∼ 5-fold reduction), rather than in ciliated cells (∼ 3.5-fold reduction) (**Figure 3A**). In the presence of IFN λ1 alone, IAV transcripts were only detected in ciliated cells indicating, similarly to 6’SLN-CD, a greater reduction of viral replication in the non-ciliated compartment compared to the ciliated one (**Figure 3A**). Almost no viral reads were detected in the 6’SLN-CD + IFN λ1 condition, independently of the epithelial cell types (**Figure 3A**). Accordingly, in the presence of both treatments the number of infected cells measured by FACS accounted for less than 1% of the total epithelium (**Figure S3**). These results further confirmed the synergistic action of IFN λ1 and 6’SLN-CD and shed light on the cell type-specific effects of the treatments. Interestingly, in presence of 6’SLN-CD alone viral replication was hindered preferentially in secretory rather than in ciliated cells. This difference was probably determined by both IAV receptor specificity and the higher susceptibility to the infection of ciliated cells, which explains the stronger reduction of viral replication observed in secretory cells. A similar explanation is also plausible for the different extents of viral replication measured across ciliated and secretory cells in the condition IFN λ1 alone. We asked whether secretory cells mounted a stronger immune response compared to ciliated cells and measured, in both cell types, the average mean expression of several key ISGs, *OAS*, *MX1*, *MX2*, *IFIT1*, *IFIT2*, *ISG15* and *ISG20* (*20*), across different experimental conditions. We did not observe significant differences, nor in the basal gene expression levels, nor in the induction upon infection or IFN λ1 stimulation (data not shown).

Lastly, we investigated the expression levels of IAV mRNA segments and we observed the following viral mRNA segment ratio: NEP > M2 > HA ∼ NP > NA > M1 ∼ NS1 > PA ∼ PB1 ∼ P2 (**Figure 3B**). The fractions of individual viral genes did not change across the treatments (**Figure 3B**), nor across epithelial cell types (data not shown). The spliced transcripts (M2 and NEP) had higher expression level compared to the unspliced transcripts (M1 and NS1). This finding is in line with previous reports showing that the expression of both M2 and NEP is more biased toward the later stages of viral replication, such as 48h hpi (*64, 71*).

### 2.4 scRNA-seq analysis reveals cell type specific responses to the infection and to the treatments within each HAE cell cluster

We then sighted to further investigate the heterogeneous cell responses to IAV infection and to the treatments within each epithelial cell cluster. As they represent a continuum of differentiation, the “basal differentiating into secretory” and “secretory” clusters were merged. Individual clustering (*54*) was performed analyzing each main HAE cell type independently of the others, and led to the identification of several subpopulations, or subclusters.

Basal cells were distributed across six subclusters annotated as follows: b1) steady state basal cells; b2) & b3) *LY6D*+ differentiating cells (*55*), with b3 displaying a more pronounced expression of *KRT14* and *KRT16*, markers of tissue repair and regeneration (*72, 73*); b4) highly proliferating cells, based on strong expression levels of genes involved in cell cycle progression such as *MIKI67*, *CDK1* and *BIRC5*; b5) proliferating cells with lower levels of cell cycle progression genes, compared to b4, but with high levels of *KRT14* ^high^ and lastly b6) inflamed cells, based on high expression of *CXCL10*, *CXCL11* and several others ISGs (**Figure 4A** and **S8A**). The latter subcluster was the less represented in the mock steady state control, while it became the most abundant in the infected untreated condition (16-fold increase), with an inflammation signature stronger than that induced in the other subclusters (**Figure 4A and S8A**). IAV induced the expression of pro-inflammatory cytokines across all basal subpopulations. 6’SLN-CD and IFN λ1, administered alone or in combination, counteracted this effect (**Figure S8A**). Similarly, the b6 inflamed cluster was decreased by 3-fold by the individual treatments and by 9-fold by the combined formulation. In turn, the levels of differentiating basal clusters (b2 and b3), which were decreased by the infection (2-fold and 3-fold decrease for b2 and b3, respectively) were also restored by the antivirals. In line with previous reports (*74*), IAV infection also reduced the b4 highly proliferating subcluster (2.8-fold decrease), which was not recovered by the individual, nor by the combined treatments (**Figure 4A**). This may also result from the inflammatory response triggered by IFN λ1, as b4 is less abundant also in the mock double treated, compared to the mock steady state (2-fold decrease). On the other hand, the b5 low proliferating cells cluster did not undergo significant changes across the tested experimental conditions.

**Figure 4.**
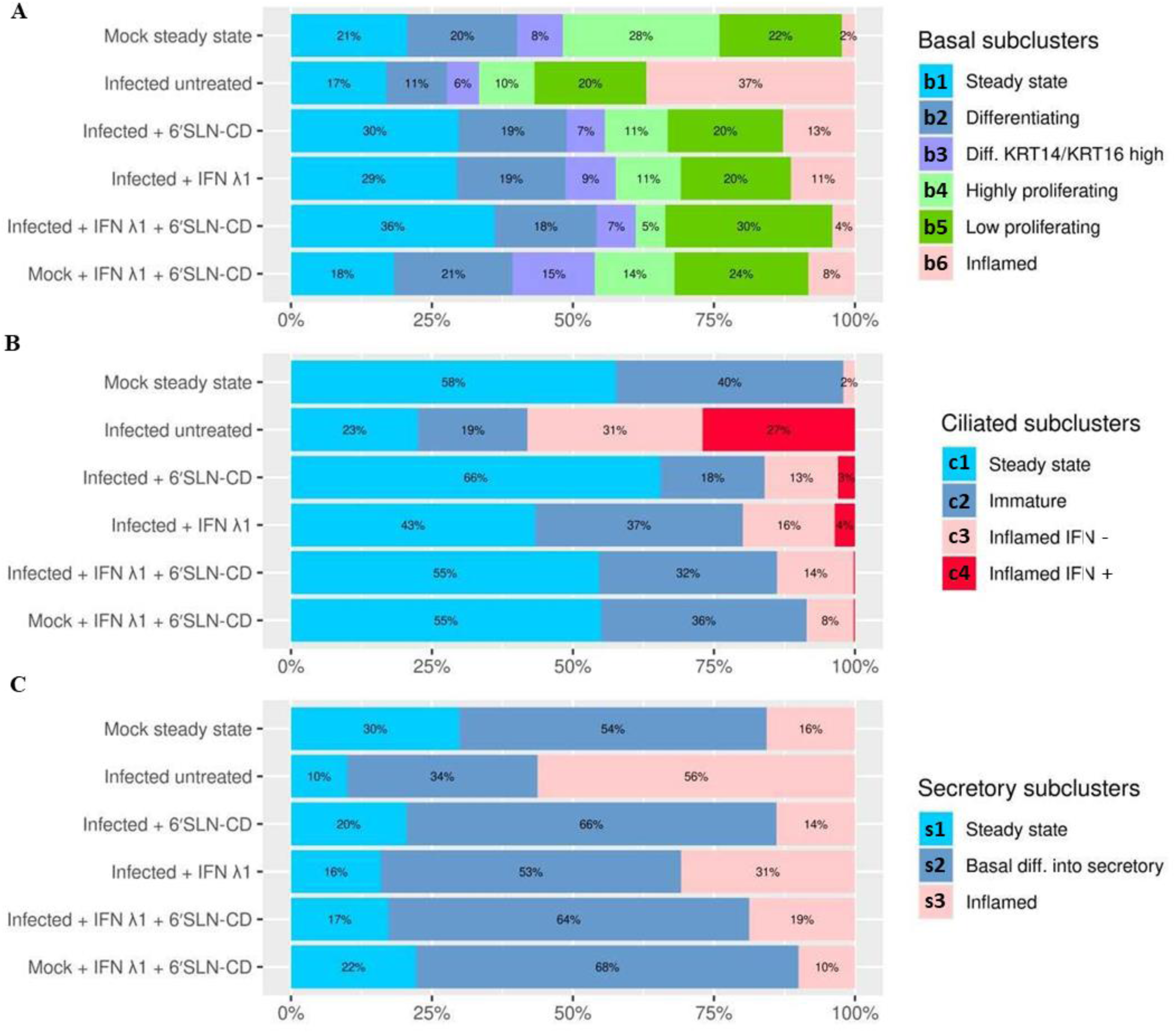
Stacked bar graph showing the relative percentages of HAE basal **A)**, ciliated **B)** and secretory cells **C)** subclusters across experimental conditions. “Diff.” stands for “differentiating”. Conditions as specified in figure 2C.

Of note, as they are not a direct target of IAV (**Figure 3A** and **S6**), basal cells mainly contributed to the immune reaction against the virus as bystander cells (*75*). Thus, all the changes induced by IAV in this epithelial compartment were largely independent of viral replication.

Within the ciliated compartment we identified four subclusters: c1) steady state cells with high expression of ciliated hallmark genes *FOXJ1*, *TPPP3*, and *ERICH3*; c2) immature cells, based on lower levels of the ciliated hallmark genes, and on higher expression of *RAB11FIP1*, which is involved in primary ciliogenesis (*76*); c3) *IFN -* inflamed cells and c4) *IFN +* inflamed cells, both characterized by high expression of inflammatory genes, such as *ISG20* and *GBP1*, but differing from each other based on the expression of *IFN λ* and *IFN β1* (**Figure 4B** and **S8B**).

Ciliated cells are highly permissive to IAV infection (*63*). Analysing the distribution of viral transcripts, we found that viral replication occurred across all ciliated subclusters (**Figure S8B**). However, c4 displayed the highest levels of IAV segments, resulting in 100% of infected cells (**Figure S8B**). The viral load correlated with the entity of the inflammatory response, as only c4 expressed *IFN λ* and *IFN β1* genes, as well as high levels of *NEDD9*, which is associated with IAV-induced antiviral response (*77*). Moreover, compared to all other clusters, c4 exhibited high levels of the pro-apoptotic factor *BBC3,* and lower or null levels of ciliated hallmark genes, probably as a result of the massive viral genes expression, hijacking the host transcriptional machinery (*46*) (**Figure S8B**). The changes in the proportions of ciliated cells subclusters across experimental conditions reflected the efficacy of the treatments. Compared to the mock steady state control, IAV infection resulted in a relative decrease of both the steady state and immature cells (2.6-fold reduction of c1 and 2.1-fold reduction of c2, respectively), whose levels were restored by both individual and combined treatments (**Figure 4C**). In line with that, 6’SLN-CD and IFN λ 1 counteracted the increase in the c3 inflamed IFN- subcluster (**Figure 4C**). The C4 inflamed IFN+ subpopulation followed a similar trend, but completely disappeared in the 6’SLN-CD + IFN λ 1 condition, further proving the combinatorial effect of the two compounds. Interestingly in the mock treated condition, the inflamed IFN - cluster was increased compared to the mock steady state but not the inflamed INF +, indicating that the latter represents a virus-specific signature at 48 hpi (**Figure S8B**).

Secretory cells were classified in three subclusters: s1) steady state secretory cells comprising a mixed population (defined as “mixed” because the gene expression profiles did not allow unambiguous classification) of mainly club cells and fewer goblet cells, displaying high expression of *SCGB1A1*, *SCGB3A1*, *MUC5AC*, *RARRES1* and *LCN2*; s2) *SCGB3A1-/TP63-* basal differentiating into secretory cells, initially described in Figure 2), expressing both secretory markers, such as *BPIFB1*, *ATP12A* and basal markers, like *KRT5*, *KRT17* and *CYP1B1*, which is exclusive of basal differentiating cells together with *LY6D* (*55*)) and s3) secretory cells differing from s2) based on lower expression of *RARRES1* and higher expression levels of ISGs (**Figure 4C** and **S8C**).

IAV infection triggered the expression of pro-inflammatory cytokines in all secretory subpopulations, however this effect was stronger in the s3 subcluster, whose fraction was increased by 3.6-fold, at the expenses of the others (**Figure 4C** and **S8C**). Secretory cells represent the second target of IAV after ciliated cells. Viral reads distribution analysis across subclusters showed that IAV preferentially infected mature rather than basal differentiating into secretory cells (**Figure 3A** and **S8C**). We did not observe an additive effect of the treatments in secretory cells: compared to the infected untreated condition, IFN λ1 alone decreased the faction of s3 inflamed cells by only 1.8-fold, while 6’SLN-CD alone restored the secretory subclusters composition as effectively as the combined treatments. This is probably due to the fact that similarly to basal cells, most of the changes occurring in secretory cells after IAV infection were largely independent of viral replication.

Our findings show that IAV infection alters the subclusters composition in epithelial cell type by inducing the appearance of inflamed populations. As the inflammatory response tightly correlates with the viral load, the ability of the antiviral treatments to restore the tissue composition to the steady state level is stronger in infected rather than in bystander epithelial cell types.

## 3. Discussion

Influenza can be a dreadful disease, with a strong socio-economic impact worldwide. Current antiviral strategies are only efficient at the early stages of the infection and are challenged by the genomic instability of the virus. We are in need for novel antiviral therapies targeting the respiratory immune defense to improve viral clearance, reduce the risk of bacterial super-infection, and attenuate tissue injury. A treatment that would prevent viral entry and at the same time boost the host antiviral response without causing immunopathology would thus represent an ideal tool to prevent or treat influenza infection. With this in mind, we assessed the antiviral potential of co-administering human IFN λ1 with 6’SLN-CD against H1N1 IAV in *ex vivo* HAE. The IFN λ1 + 6’SLN-CD formulation is non-toxic and more effective in reducing viral replication, compared to the individual treatments. IFN λ1 has been already used in clinical trials against viral infections, while 6’SLN-CD is well tolerated *in vivo* and effectively constrains the spread of IV infection when administered topically (*36*). Overall, our data support a prospective therapeutic application of IFN λ1 + 6’SLN-CD.

We next sought to investigate the mechanism of action of this formulation by scRNA-seq in HAE. Transcriptomic analysis unraveled the heterogeneous composition of each main epithelial cell type, which is an assortment of subclusters with unique gene expression programs underlying different cell states. Besides terminally differentiated cells we also identified basal differentiating into secretory cells. This subpopulation, roughly equally represented across experimental conditions, derived from the continuous differentiation process occurring in the respiratory epithelium. We did not find basal cells differentiating directly into ciliated cells, a process triggered by tissue injury (*58, 78, 79*), because in our settings the cytopathic effect induced by IAV at 48 hpi is not strong enough to significantly alter the architecture of the tissues (*11*). We also did not observe secretory cells differentiating into ciliated cells ^[54-56]^, probably due to the limitations in sequencing dept and to the fact that we did not perform a lineage study (*57*), which would be beyond the scope of this work. When we measured the distribution of viral reads across the main cell types, we found that IAV preferentially infected epithelial cells in the following order: ciliated, secretory and basal cells. Accordingly, ciliated cells mounted a stronger inflammatory response compared to secretory cells which, in turn, expressed ISGs and innate cytokines at higher levels than basal cells. Interestingly, only within the ciliated compartment, IAV induced the appearance of highly inflamed cells characterized by a distinctive high expression of IFN type I and type III genes. Basal cells, which were the most diverse due to their multipotent potential, displayed extremely low levels of viral transcripts and participated to the tissue immune response as bystander cells. Of note, the infection of basal cells would be highly detrimental to the host as these cells are absolutely necessary to maintain the barrier of the respiratory epithelium by regenerating secretory and ciliated cells targeted by IV (*79*). We observed that the proportions of the main HAE cell type clusters did not change upon infection and/or treatments which is, as explained above, due to the poor cytopathic effect induced by IAV at 48 hpi (*11*). On the other hand, the subcluster composition of each cell type underwent significant modifications in response to the infection, mostly resulting from the appearance of inflamed cells. These changes were induced in ciliated and to some extent in secretory cells, by a direct cell response to viral replication and in basal cells by the response to the paracrine signaling from infected cells.

We observed that the individual treatments reduced the percentage of viral transcripts to different extents across epithelial cell types: both 6’SLN-CD and IFN λ1 alone caused a reduction of viral reads more pronounced in secretory rather than in ciliated cells. This finding was unexpected for 6’SLN-CD, which is designed to exclusively target extracellular viral particles and was evenly distributed throughout the apical side of the HAE. We reasoned that the effect of the 6’SLN-CD in reducing viral replication depends on the susceptibility of the cells to the infection, which is dictated by both the distribution of α 2,6-linked Sia and the expression of host factors necessary for viral entry. As IAV infects ciliated cells more easily than secretory cells, the number of viral transcripts is lower in the latter cell type and in turn, its reduction in response to 6’SLN-CD is more pronounced, compared to that observed in ciliated cells.

Viral transcript distribution analysis also indicated a synergistic effect between 6’SLN-CD and IFN λ1 in each main cell type. However, the capacity of the combined treatments to revert IAV-induced perturbations in subclusters composition was greater than the individual ones only in ciliated cells, where the inflammation was a direct consequence of viral replication and in basal cells, but only limited to the inflamed subcluster. In line with that, in the basal compartment where changes in cell composition were independent of IAV infection, nor the individual nor the combined treatments succeeded in restoring the number of highly proliferating cells. Lastly, in absence of infection, the combined treatment did not alter the ratios between the main basal, ciliated and secretory cells clusters, but changed the subclusters abundances within each of them, resulting in an increase in inflammatory cells which was likely induced by IFN λ1.

Different macromolecules-based approaches are currently available for the treatment of viral infections. However, a deep knowledge of the impact on the host cells is needed to increase the effectiveness of these therapies, minimize the side effects and reduce the toxicity. Our study, proposing scRNA-seq to assess the effects of a combined therapy against IV, is in line with this need and to the best of our knowledge is the first to present such analytical approach. We suggest that the ability of an antiviral treatment to restore epithelial cell subclusters composition upon infection is an important parameter to dissect the effects of the drug on the host cells, beyond its capacity to impair viral replication. This work is also the first in addressing at the molecular level the anti-IAV effects of IFN λ in HAE. Additional investigations in more relevant *in vivo* models of infection, such as mice or ferrets, will be necessary to further assess the efficacy of IFN λ1 + 6’SLN-CD formulation and its genetic barrier to antiviral resistance. Also, in light of the ongoing differentiation process occurring in the respiratory epithelium, scRNA-seq velocity analysis (*80*) could allow to investigate the trajectories of both basal and secretory cells differentiation, as well as how such trajectories would be perturbed by the infection and the treatments.

## 5. Materials and Methods

*Human airway epithelia (MucilAir™):* the human airway epithelia used in this study were reconstituted from freshly isolated primary human nasal polyp epithelium collected either from 14 different donors, upon surgical nasal polypectomy, either from individual donors, as previously described (*11*). The patients presented with nasal polyps but were otherwise healthy, with no atopy, asthma, allergy or other known comorbidity. All experimental procedures were explained in full, and all subjects provided written informed consent. The study was conducted according to the Declaration of Helsinki on biomedical research (Hong Kong amendment, 1989), and the research protocol was approved by the local ethics committee (*11*). The tissues were maintained at the air-liquid interface according to the manufacturer’s instructions (*11*).

*Viral stocks and compound*: influenza H1N1 A/Switzerland/3076/16 clinical specimen was isolated from the nasopharyngeal swab of an anonymized patient, provided from the Geneva University Hospital. The sample was screened by one-step real-time quantitative PCR (qPCR) (*81*). Influenza A virus was subtyped by sequencing the NA gene as previously described (*82*). To prepare viral stocks, 100 µl of clinical sample was inoculated at the apical surface of several HAE, for 4h at 33°C. After the infection the apical side of the tissues were washed five times with PBS. In order to measure the daily viral production, apical samples were collected every 24 h, by applying 200µl of medium for 20’ at 33°C. The viral load of each time point was then measured by RT-qPCR and the 4 apical washes with highest titer were pooled and re-quantified. Aliquots were stored at -80°C.

Human recombinant IFN λ1 protein was obtained from R&D Systems, Inc. (Abingdon, United Kingdom). 6’SLN-CD was synthetized as described previously (*36*).

*HAE, viral infections and treatments*: HAE were infected apically with H1N1 A/Switzerland/3076/16 strain (1e3 or 1e4 RNA copies/tissue), in a final volume of 100 µl as described above (*83*). Infected tissues were treated with 6’SLN-CD alone, IFN λ1 alone, or with 6’SLN-CD plus IFN λ1. 6’SLN-CD dissolved in PBS was transferred on the apical surface of the tissues (60 μg/tissue, in a volume of 30ul), starting from 8 hpi. After each apical wash, performed for daily viral load quantification as described above, 6’SLN-CD was re-added on the apical side of the tissues. IFN λ1 was added on the basal side of the inserts (5.5 ng/tissue in 550 µl) at one day before infection and then added every day, each time replacing the entire basal medium with fresh one. In parallel, upon each wash the infected untreated tissues were administered with 30 µl of PBS on the apical side, the same volume added to the tissues treated with 6’SLN-CD, while the basal medium was changed on a daily basis. The treatments were administered up to 72 hpi.

*Viral load quantification*: viral RNA was extracted from Mucilair apical washes using EZNA viral extraction kit (Omega Biotek) and quantified by using RT-qPCR with the QuantiTect kit (#204443; Qiagen, Hilden, Germany) in a StepOne ABI Thermocycler, as previously described (*11*). Viral RNA copies were quantified as follows: 4 ten-fold dilution series of *in vitro* transcripts of the influenza A/California/7/2009(H1N1) M gene were used as reference standard as previously described (*11*). CT values were converted into RNA load using the slope-intercept form. In all experiments, the slope, efficiency and R2 ranged between 0.96 and 0.99 (*38, 84*). *P values* were calculated relative to untreated controls using the two-way ANOVA with Prism 8.0 (GraphPad, San Diego, CA, USA).

*Toxicity and viability assays*: non-infected tissues were treated with 6’SLN-CD plus IFN λ1, in the same doses/volumes used for infected tissues (as described above). Accordingly, every day an apical wash was performed and a new dose of 6’SLN-CD was added on the apical side of the tissues, while fresh medium with IFN λ1 was added on the basal side. Similarly, the untreated control tissues were administered with 30μl of PBS on their apical side, while the basal medium was replaced every day.

Lactate dehydrogenase (LDH) release in the apical medium was measured with the Cytotoxicity Detection Kit (Roche 04744926001) as described previously (*36*). Percentages of cytotoxicity were calculated compared to the cytotoxicity control tissues, which were treated with 100 μl of PBS-5% Triton ™ X-100 (Sigma Aldrich) on the apical side.

Cell metabolic activity was measured by adding MTT reagent (Promega), diluted in MucilAir medium (1 mg/ml), on the basal side of the tissues. After 3h at 37°C the tissues were transferred in a new plate and lysed with 1 ml of DMSO. Subsequently, the absorbance was read at 570 nm according to manufacturer instructions. Percentages of viability were calculated by comparing the absorbance to the untreated control tissues.

*HAE enzymatic dissociation and flow cytometry analysis*: HAE tissues were first washed both apically and basally in DPBS without calcium and magnesium (Thermo Fisher Scientific) for 10’ at 37°C. Then they were incubated with TrypLE (Thermo Fisher Scientific), both apically and basally for 30’ at 37°C. During this time the tissues were dissociated with a 1ml pipette. Cells were harvested and washed with ice-cold MACS buffer (PBS without calcium and magnesium, EDTA pH 8, 2mM BSA 0.5%).

For scRNA-seq, cells were stained with Hoechst 33342 (Thermo Fisher Scientific) and DRAQ7 (Biolegend) and analyzed with a MoFlo Astrios Cell Sorter (Beckman Coulter). Viable cells were defined as Hoechst +/DRAQ7-, doublets were excluded by gating for SSC-W vs SSC and single cells were sorted.

To determine the percentages of infected cells, upon HAE dissociation the cells were fixed/ permeabilized using the Perm/Wash Buffer RUO (554723 BD Biosciences-US) and then stained with the primary antibody (mouse monoclonal anti-IVA Ab 1:100 dilution, Chemicon®) for 20’ at 4°C. After a wash with Perm/Wash Buffer RUO the secondary Ab (Alexa Fluor 488 Invitrogen™, 1 :200) was added for 20’ at 4°C. After one wash with MACS buffer the percentages of IAV infected cells were determined with a MoFlo Astrios Cell Sorter (Beckman Coulter) and the uninfected gating control was defined using uninfected cells stained with both the primary and the secondary antibodies.

*Single cell RNA sequencing of HAE*: upon HAE dissociation viable cells were sorted as described above. Cells were then counted using Countess™ II FL Automated Cell Counter (Invitrogen) and diluted to equivalent concentrations with an intended capture of 5000 cells/sample, following the manufacturer’s provided by 10x Genomics for the Chromium Single Cell platform. All subsequent steps through library preparation followed the manufacturer’s protocol. Samples were sequenced on an Illumina HiSeq 4000 machine.

*Computational analysis of scRNA-seq data*: upon demultiplexing and performing the routine quality checks on the raw reads, we processed the data via Cell Ranger version 3.1.0 using the union of human and Influenza A genome as a reference (see References and Annotations) (*85*). We extracted the UMI counts for the 10000 most frequent cell barcodes in each sample, then screened the distributions of total UMI counts, percentages of mitochondrial and viral genes across for these barcodes (within each sample), and, finally, selected cell barcodes having more than 10000 reads and not more than 15% of mitochondrial reads for downstream analysis (*86*). The analysis of single-cell data was performed using Seurat version 3.2.3 (*87*). First, the raw UMI counts were transformed to normalized expression levels on a common scale using SCTransform method implemented in Seurat which amounts to computing (Pearson) residuals in a regularized binomial regression model for UMI counts (*88*). The normalization was performed jointly on all samples and the genes expressed in less than 10 cells were excluded prior to normalization together with the viral genes.

The selected cells from all samples were first clustered on the normalized expressions of hallmark marker genes only which resulted in five stable clusters. The clustering method implemented in Seurat we applied amounts to 1) constructing a *k*-nearest neighbor graph of all cells, 2) deriving an (approximate) shared nearest-neighbors graph, and 3) applying a modularity-based community detection to the latter graph (*89*). Euclidian metric was used for the construction of *k*-NN graph.

Out of the five identified global cluster lacked a clear signature and was contained a sizeable proportion of cells with high total UMI counts compared to other clusters (Figures 3, S3 and S4). Given it included only a small percentage of cells we excluded it from further analysis hypothesizing that this cluster likely contains a large percentage of doublets. The remaining four clusters were clearly identified as basal cells, ciliated cells, secretory cells and basal cells differentiating into secretory cells (Figures 3, S4). The latter two clusters were merged for further analysis.

The cells in the identified global cell-type clusters were then analyzed in isolation from each other in order to identify cell-type specific responses. For each cell type we first found a tentative set of genes differentially expressed between conditions and then re-clustered the cells based on their expressions across these genes. The same graph-based clustering was applied with cell distances derived from the first 10 principal components of the expression matrix. Condition-differential genes were found as a union of genes overexpressed in any condition versus the rest as assessed by the Mann-Whitney-Wilcoxon test with the nominal p-value of 0.01. The obtained differential genes were ordered using a hierarchical clustering algorithm and manually curated before producing the cell-type specific heatmaps shown in Figure S7.

To test for the differential expression of host factors between secretory and ciliated cells in steady state conditions we first constructed a list of 52 host factor genes and retained those expressed in more than 50% of secretory and ciliated cells in (steady state condition) which resulted in 33 genes. We then tested for the differences in normalized expression between secretory and ciliated cells using Mann-Whitney-Wilcoxon test (with a two-sided alternative) and adjusted the resulting p-values using Bonferroni correction. Genes displayed in Figure S7 were selected manually.

*Human genome annotation:* GRCh38.p10 with only the main chromosome contigs retained i.e. chr1-chr22, chrX, chrY and chrM. *Human genome annotations:* Gencode release 29 with annotations of non-gene features (e.g. exons, transcripts, CDSs and UTRs) removed if they overlapped with protein-coding or lincRNA features and did not have “protein-coding”, “lincRNA” or “processed-transcript” tags themselves. *Influenza A reference and annotations:* GCA_001343785.1 (*90*). The viral reference annotations were preprocessed by prefixing all gene ids by “InfluenzaA” and by changing the type of “CDS” features to “exon”.

## Acknowledgments

We would like to thank the Genomic Platform at the University of Geneva) and Dr. Christel Borel for providing precious assistance and support in data analysis and experimental design. This work was supported by the Swiss National Science Foundation (Sinergia grant CRSII5_180323 to F.S. and C.T.) and by the Fondation Aclon (Geneva, to C.T.).

## Conflict of interest

The authors declare no conflict of interest. The funders had no role in the design of the study; in the collection, analyses, or interpretation of data; in the writing of the manuscript; or in the decision to publish the results.

## Author contribution

C.M. and I. K. Contribute dequally to this work. All authors designed research. C.M., Y.Z., S.C., S. H., A.C-A.Z. and V.C., performed research. C.M. and I. K., analyzed data. C.M. and C.T. wrote the paper.

## Supporting Information

**Figure S1.**
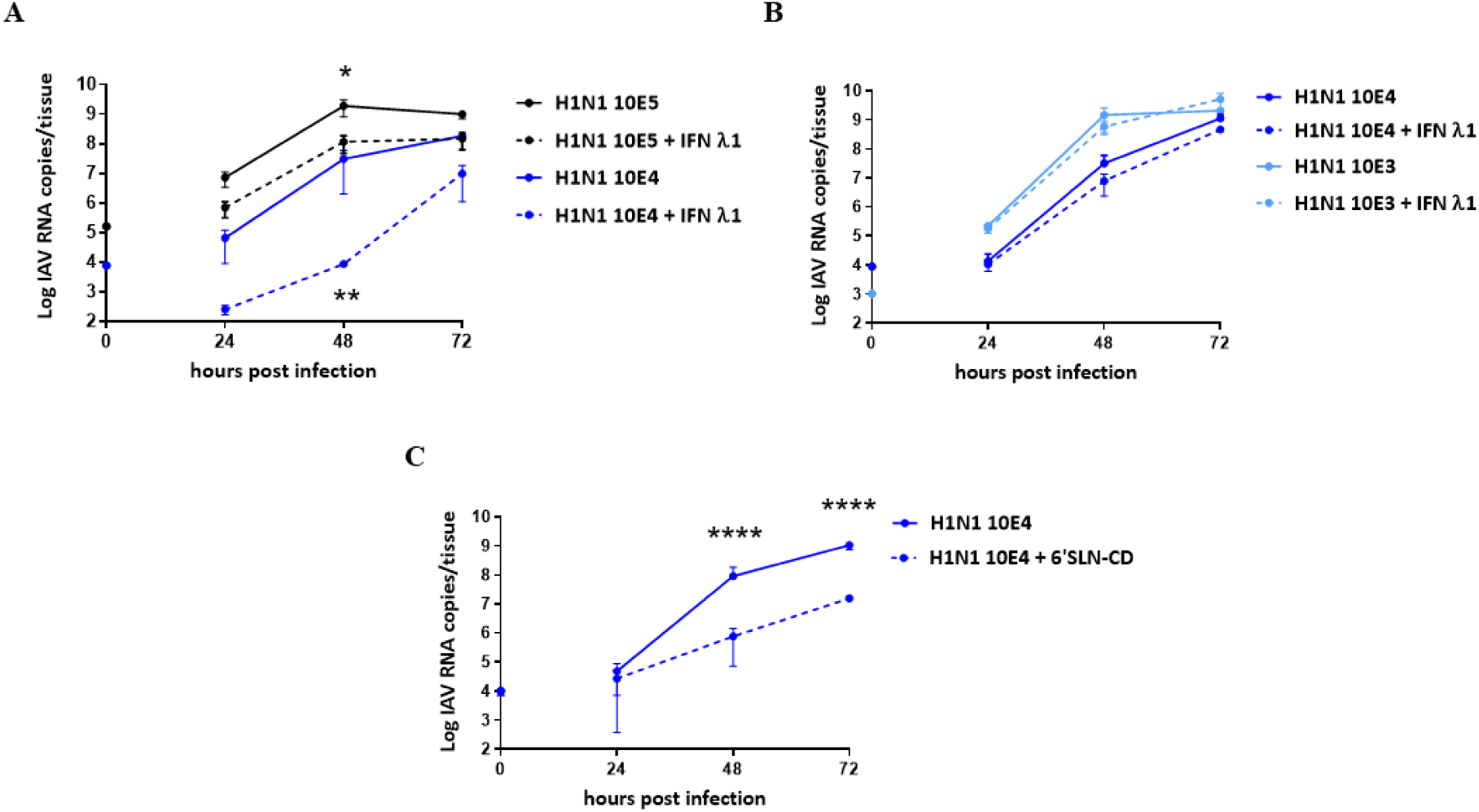
Antiviral activity of IFN λ1 and 6’SLN-CD in HAE. HAE developed from a single donor were treated on their basal side with IFN λ1 (5.5 ng/insert) either starting at 24 hbi **A)**, or at 8 hpi **B)**, and infected with different numbers of RNA copies of clinical A/Switzerland/3076/2016 H1N1 (0 h corresponds to the time of viral inoculation). IFN λ1was then administered daily up to 48 hpi. **C)** 6’SLN-CDs (60 µg/insert) were administered daily on the apical side of the tissues, starting from 8 hpi and up to 48 hpi. Viral replication was assessed measuring the apical release of IAV by RT-qPCR. The results were obtained using HAE developed from different donors and represent the mean and standard deviation from two independent experiments. *, *p* ≤ 0.5; **, *p* ≤ 0.01; ****, *p* ≤ 0.0001.

**Figure S2.**
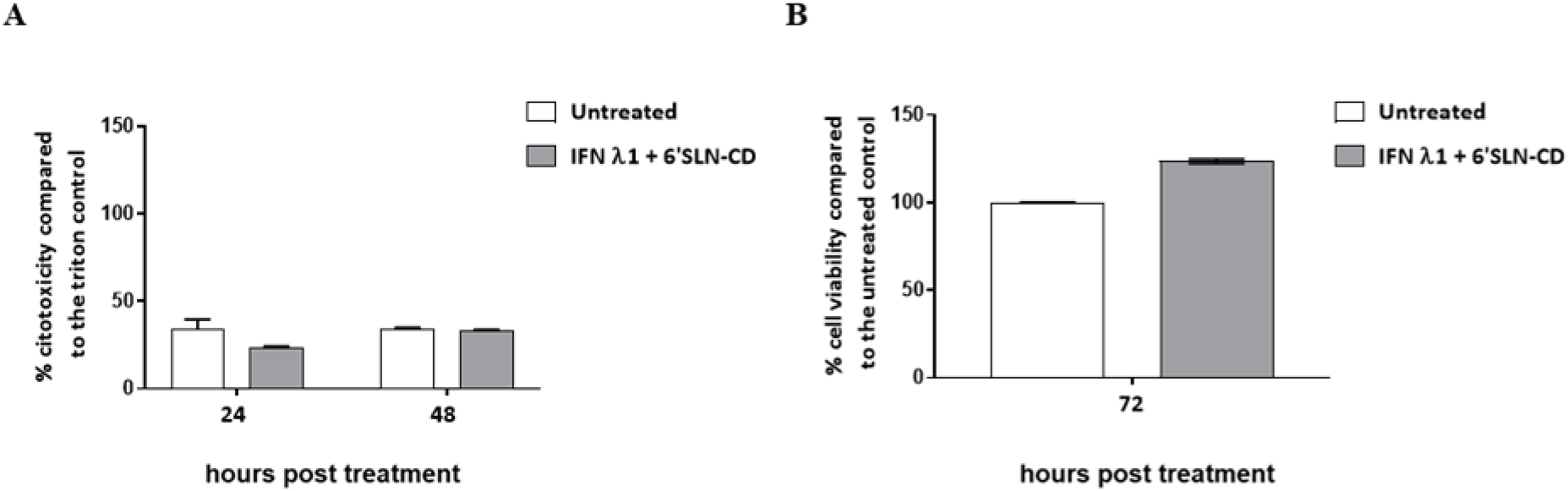
Toxicity assessment of IFN λ1 + 6’SLN-CD in uninfected HAE. **A)** Measurement of cellular cytotoxicity by LDH assay. The percentage of LDH release was calculated compared to the triton cytotoxicity control. **B)** Measurement of cell metabolic activity by MTT assay. The percentage of MTT reduction into formazan was calculated relatively to the untreated control. The tissues were treated daily with 6’SLN-CD on their apical surface and with IFN λ on their basal side (60 μg and 5.5 ng per tissue, respectively, in PBS), for 72 h. The MTT assay was performed at 72 hours post treatment (hpt), while the LDH assay was performed at both 24 and 48 hpt on the apical sides of the tissues. Untreated control tissues (untreated) and cytotoxicity control tissues were treated on their apical side with PSB or with PBS-Triton 5%, respectively. The results represent the mean and standard deviation from two independent experiments performed in duplicate.

**Figure S3.**
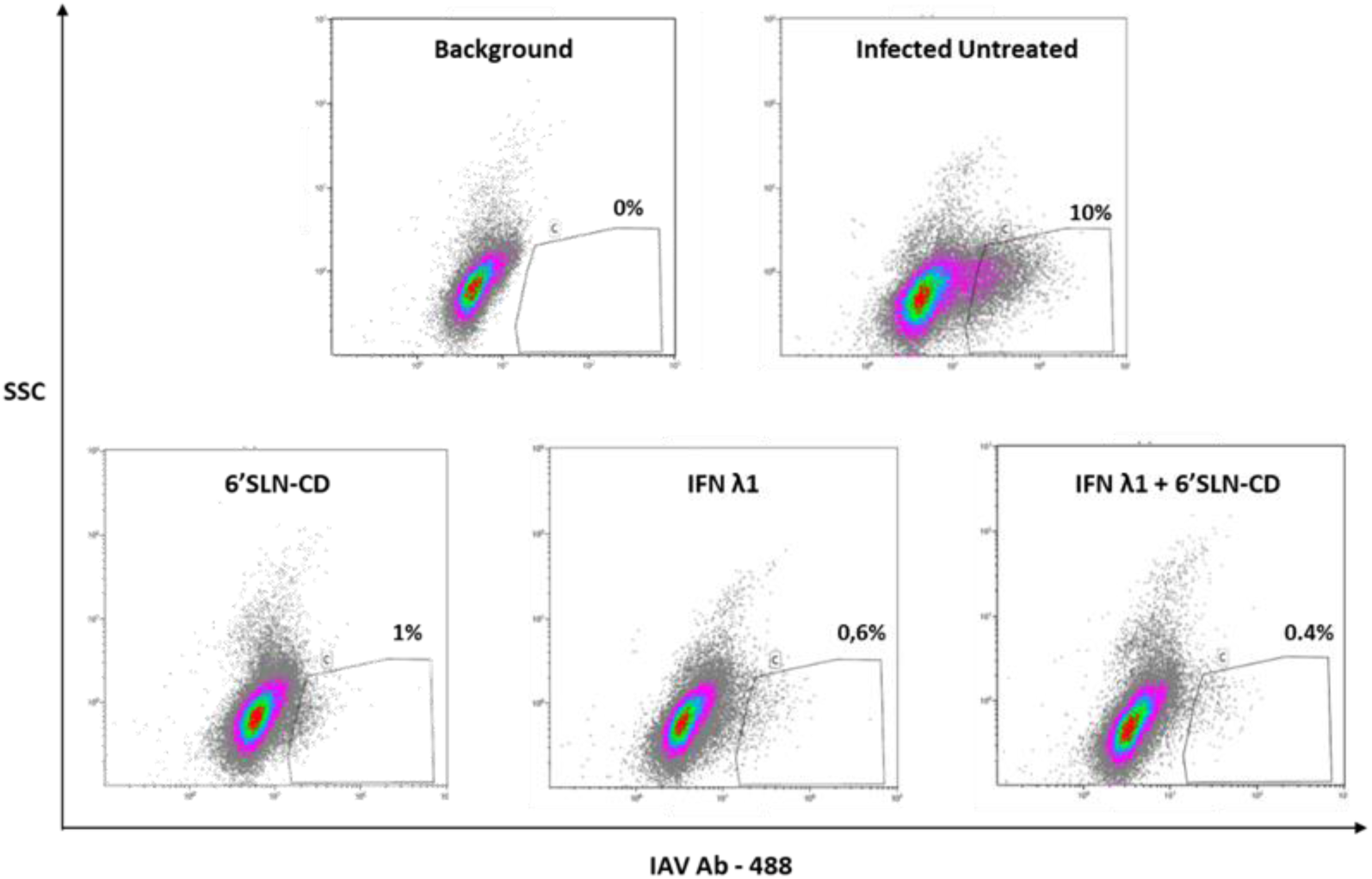
Flow cytometry (FACS) analysis of infected HAE cells at 48 hpi, with an antibody (Ab) targeting IAV nucleoprotein. The results were obtained from the same tissues shown in figure 1B and are representative of two independent experiments. HAE developed from a pool of donors were infected with 10^3^ RNA copies of clinical A/Switzerland/3076/2016 H1N1 and treated or not with 6’SLN-CDs (60 µg/tissue, administered daily starting at 8 hpi), or with IFN λ (5,5 ng/tissue, administered daily staring from 24h before infection), or with both compounds. At 48 hpi the tissues underwent enzymatic digestion and staining. The gating was defined based on the background signal obtained from an infected tissue stained with the 2^nd^ Ab alone.

**Figure S4.**
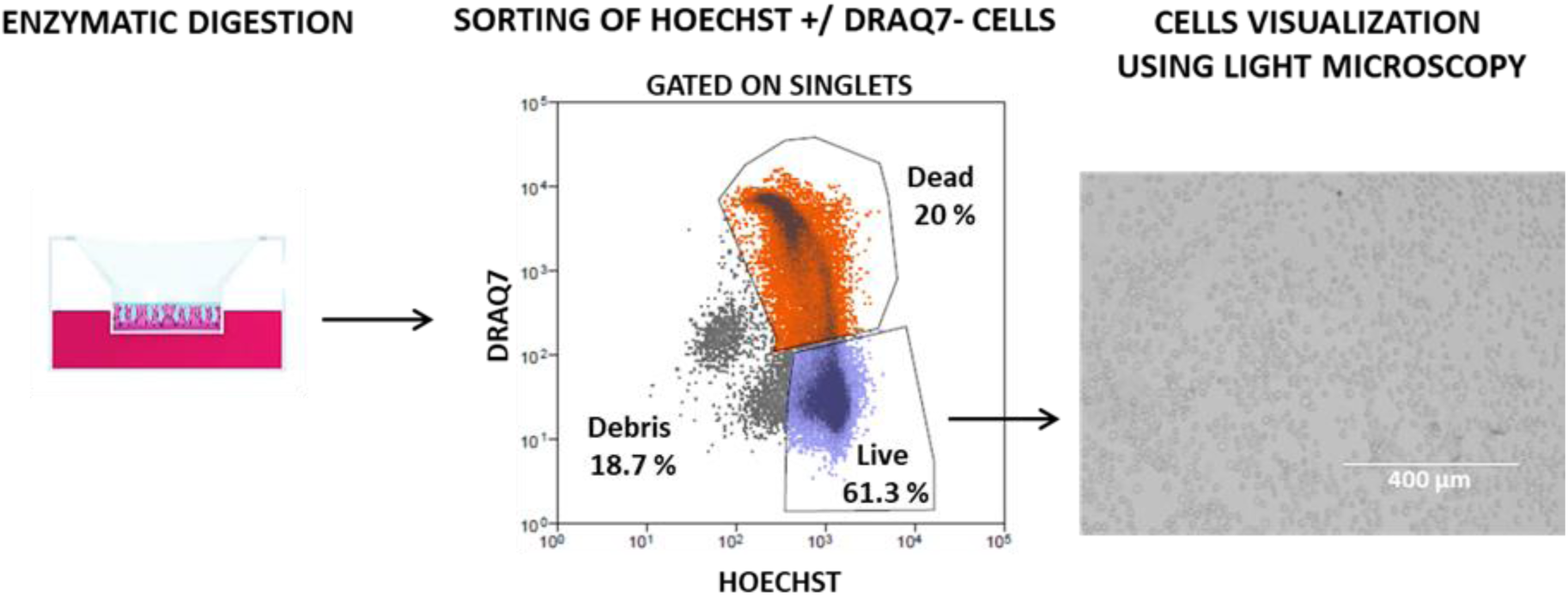
HAE dissociation protocol for scRNA sequencing. Upon enzymatic digestion, cells were stained with Hoechst to label the nuclei and with DRAQ7 to exclude non-viable cells. Hoechst +/DRAQ7 – cells were sorted, visualized at the light microscope and then processed for scRNA-seq analysis.

**Figure S5.**
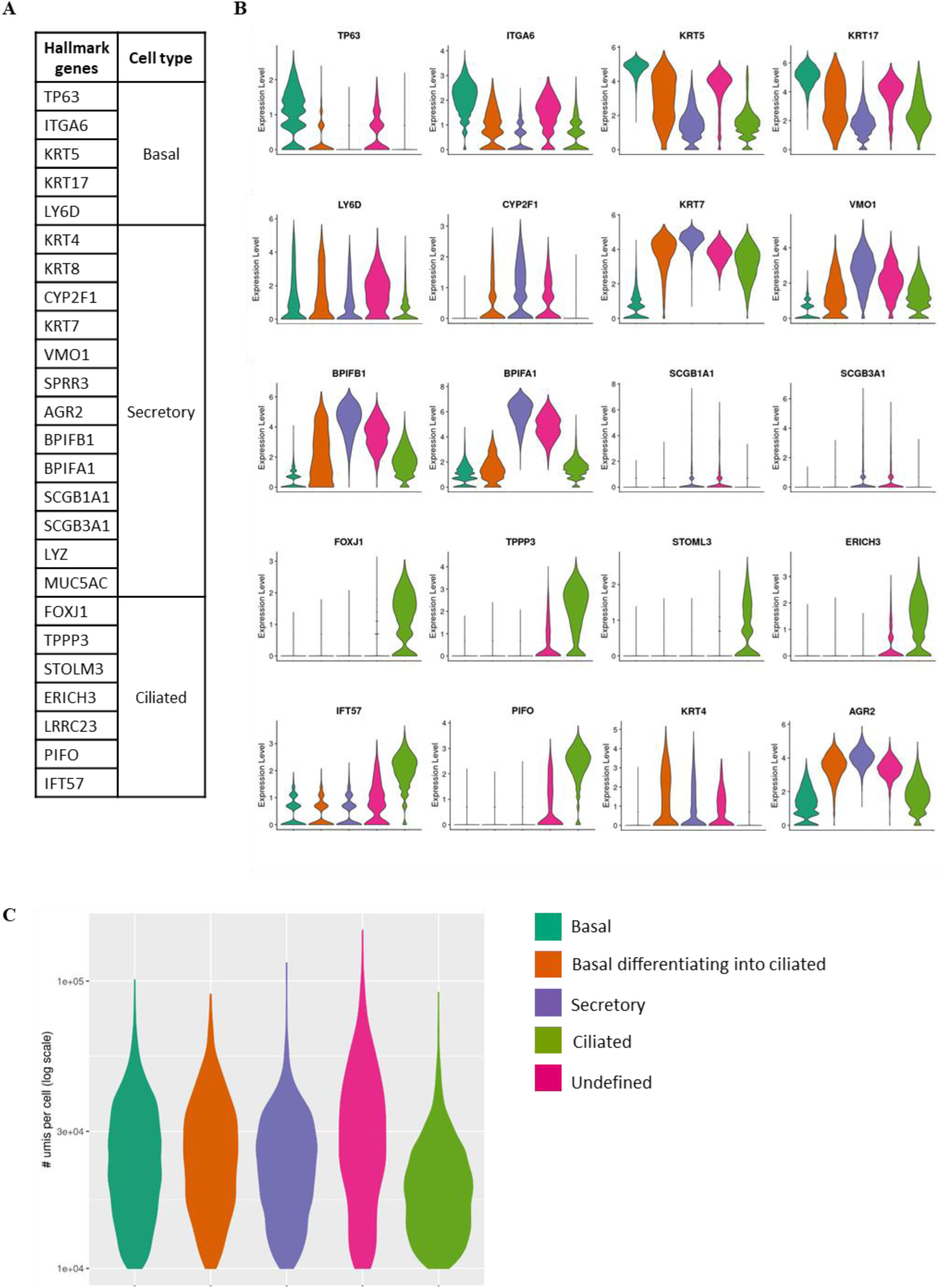
A) Hallmark genes used to annotate the main human epithelial respiratory cell types. **B)** Violin plots showing the expression distribution of cell-type specific hallmark genes across the HAE cells clusters described in Figure 2D. **C)** Violin plots showing the UMI (unique molecular identifiers) distributions across HAE cell clusters.

**Figure S6.**
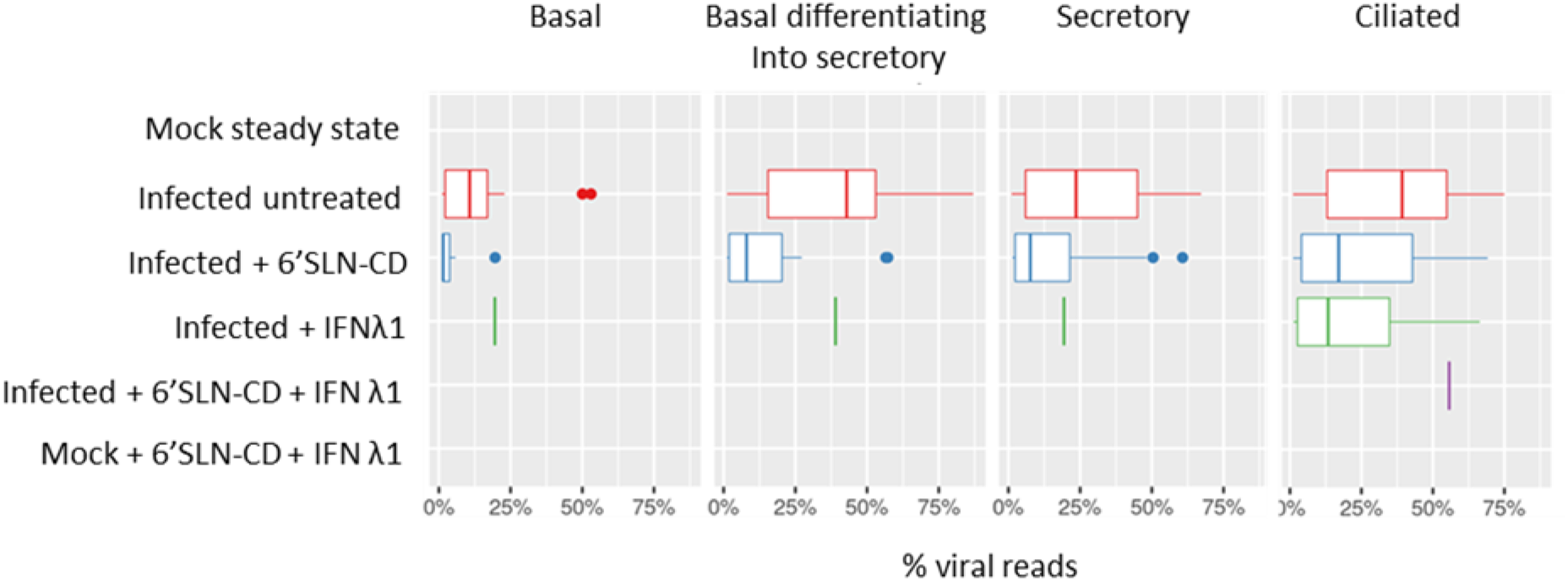
Box plot showing the distribution of viral mRNA molecules in cells having more than 1% of viral reads, across different HAE clusters experimental conditions.

**Figure S7.**
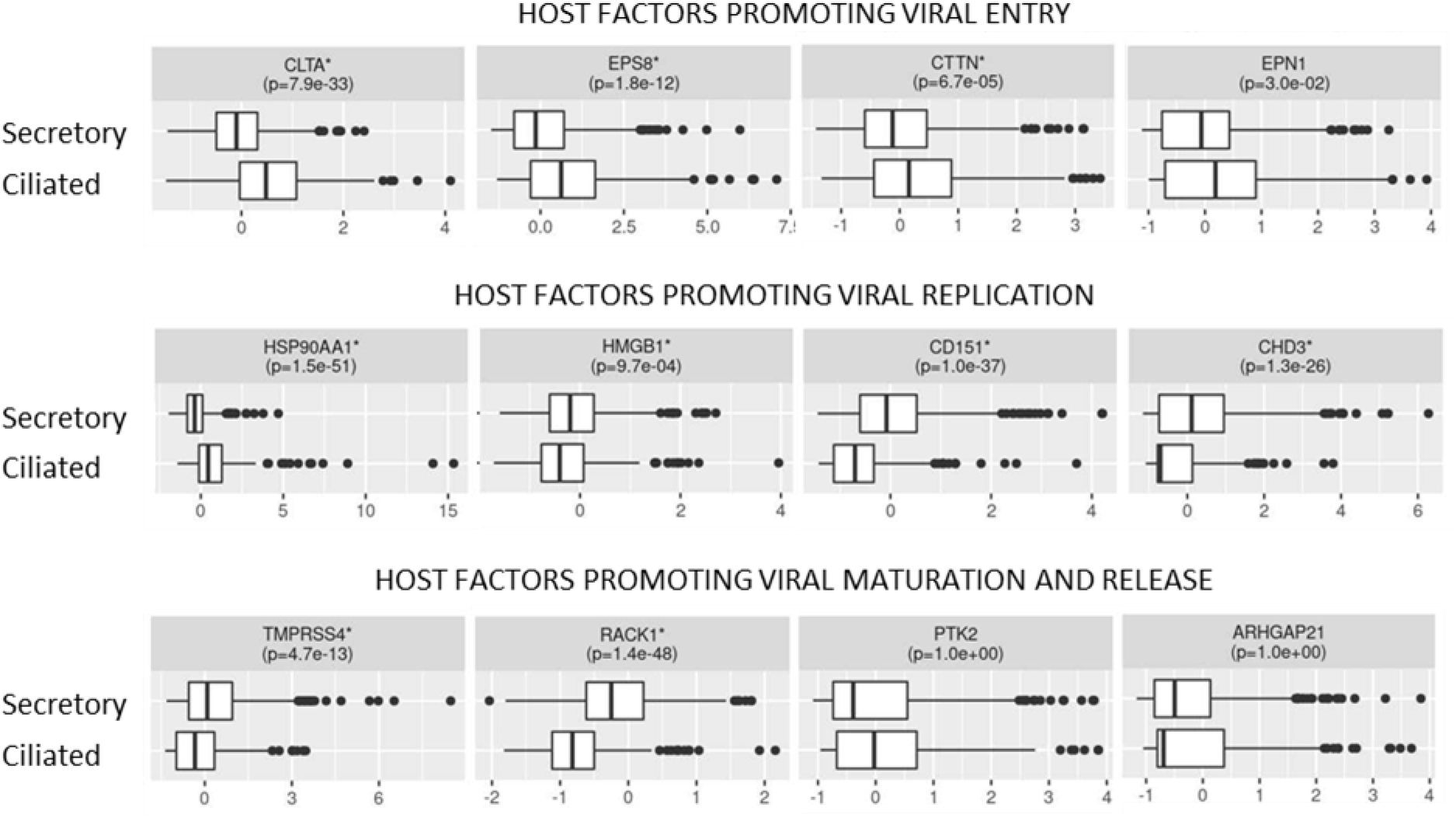
Expression of host factors involved in IV replication across secretory and ciliated cells in steady state conditions. Expression values are Pearson residuals from SCTransform binomial regression model and p-values are from Mann-Whitney-Wilcoxon test, additionally adjusted for multiple testing (see Methods).

**Figure S8.**
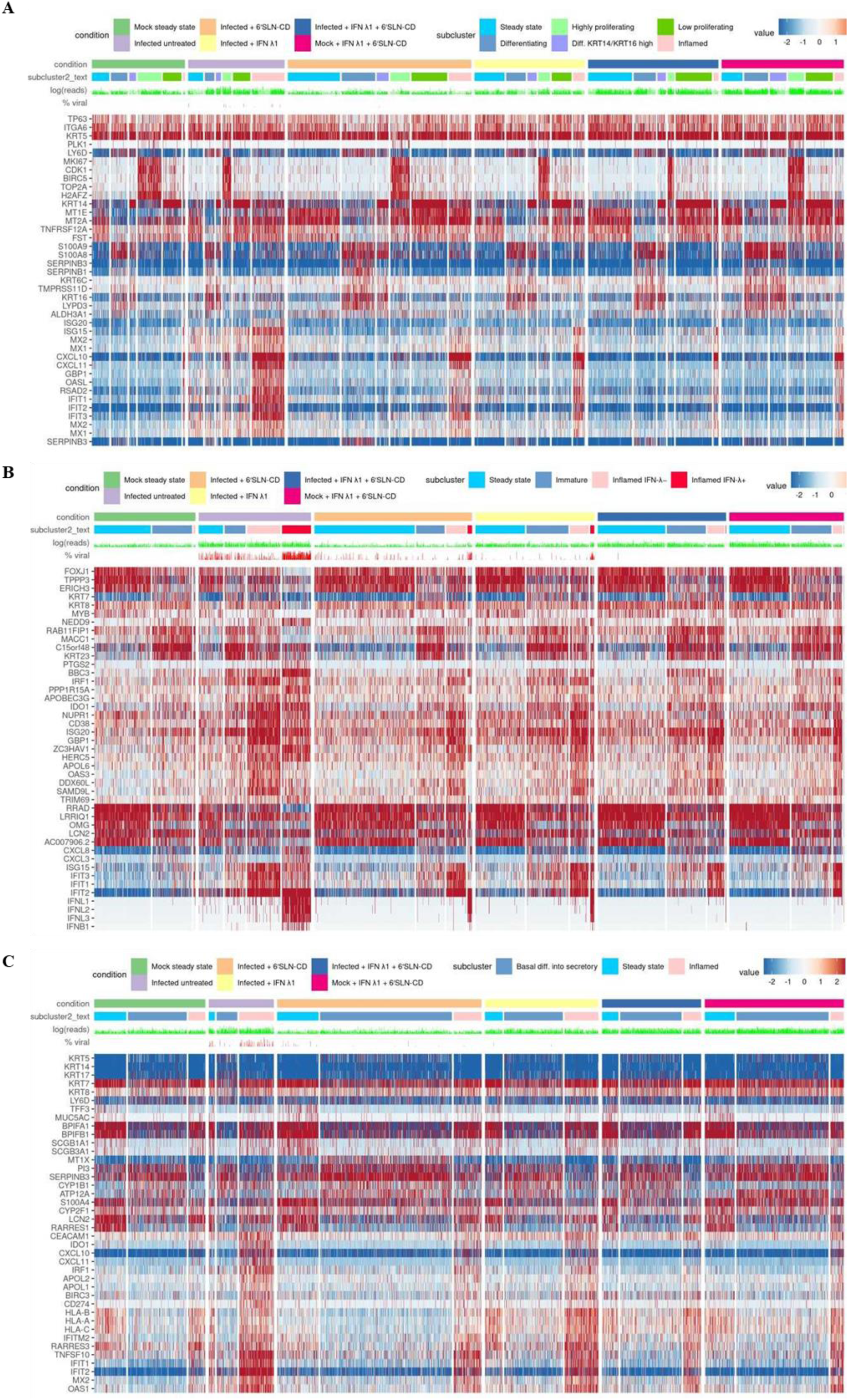
Gene expression profiles of basal (**A**), ciliated (**B**) and secretory (**C**) cells subclusters across experimental conditions (see Figure 4). The percentages of viral reads across cells are shown in red above the heatmaps, while the number of total UMI counts is shown in light green. Expression values are Pearson residuals from SCTransform binomial regression model [70] fitted to UMI counts (see Methods).

## Notes

### Competing Interest Statement

The authors have declared no competing interest.

